# A coherent feed-forward loop fine-tunes KatG to maintain redox homeostasis in mycobacteria

**DOI:** 10.64898/2025.12.17.694866

**Authors:** Xinfeng Li, Yazheng Huang, Fang Chen, Jinfeng Xiao, Xiaoyu Liu, Mingyue Zhong, Xinyu Tao, Hang Yang, Jin He

## Abstract

Bacteria survive and reproduce in fluctuating environments by balancing their basic physiological needs, such as relying on iron-containing enzymes to detoxify reactive oxygen species (ROS) while conserving scarce iron. Understanding the regulatory networks that balance these conflicting needs is key to deciphering bacterial adaptability. Here, we identify a coherent feed-forward loop in mycobacteria that integrates the transcriptional regulator FurA3, the small RNA MrsI, RNA sponge FutR, and the catalase-peroxidase KatG. This circuit fine-tunes antioxidant responses according to iron availability. Under iron limitation, the iron-containing enzyme KatG is dually repressed by FurA3 transcriptionally and by MrsI post-transcriptionally, which conserves iron at the cost of antioxidant capacity. However, when iron deprivation coincides with oxidative stress, the circuit triggers an emergency response: H₂O₂ inactivates FurA3, relieving transcriptional repression of *katG* and *futR*, while induced FutR sequesters MrsI to post-transcriptionally derepress *katG* expression, enabling rapid KatG synthesis for effective peroxide detoxification. Our findings reveal a tightly regulated network that allows mycobacteria to balance iron homeostasis and oxidative stress defense, providing mechanistic insights into bacterial physiological trade-offs.

## Introduction

Bacteria survive and reproduce in fluctuating environments by continuously managing fundamental physiological trade-offs (Basan et al., 2020). A classic and widespread challenge is their reliance on iron-containing enzymes, such as catalases and peroxidases, for the detoxification of reactive oxygen species (ROS), which forces them to balance oxidative defense with the need to conserve scarce iron (Andrews et al., 2003; Imlay, 2013). Among various environmental stresses, oxidative stress caused by the accumulation of ROS, such as superoxide anion (O₂•⁻), hydrogen peroxide (H₂O₂), and hydroxyl radical (HO•), is a major threat (Buchser et al., 2023; Gupta and Imlay, 2023; Imlay, 2013, 2019). These cytotoxic molecules, generated as byproducts of aerobic metabolism activity and induced by exogenous physical and chemical factors, damage key cellular components, including nucleic acids, proteins, and lipid (Henningham et al., 2015). To address this, bacteria have evolved highly coordinated enzymatic pathways for the stepwise detoxification of ROS. The process begins with superoxide dismutase (SOD), which converts superoxide into H₂O₂ (Broxton and Culotta, 2016). Iron-dependent catalases and peroxidases then reduce H₂O₂ and organic hydroperoxides, transforming them into harmless products (Barreiro et al., 2023; Sooch et al., 2014; Zamocky et al., 2008). Understanding the complex regulatory networks that orchestrate this critical trade-off between iron homeostasis and antioxidant defense is essential for unraveling bacterial adaptive strategies.

The regulation of bacterial antioxidant defenses is intricately linked to iron homeostasis and is controlled by complex transcriptional and post-transcriptional networks (Dard et al., 2024; Imlay, 2015; M et al., 2022). Key transcriptional regulators have been identified, such as OxyR (Choi et al., 2001; Marinho et al., 2014; Pedre et al., 2018), the iron-sensing Fur-family repressor PerR (Lee and Helmann, 2006; Marinho et al., 2014; Zhang et al., 2025), SoxRS (Seo et al., 2015), OhrR (Soonsanga et al., 2008), and SarZ (Chen et al., 2009), of which OxyR and PerR are the most extensively studied peroxide sensors. In Gram-negative bacteria, OxyR functions as a peroxide-sensing transcription factor, activating detoxification genes in response to H₂O₂ (Choi et al., 2001; Choudhary et al., 2024; Kim et al., 2002; Pedre et al., 2018). Specifically, OxyR detects H₂O₂ through oxidation of a conserved cysteine, which forms a disulfide bond with a nearby cysteine, leading to a conformational change that activates OxyR and triggers the transcription of peroxide detoxification genes. In contrast, Gram-positive bacteria primarily rely on PerR, a metalloregulator of the Fur-family that acts as a peroxide-sensing repressor (Lee and Helmann, 2006; Zhang et al., 2025). PerR senses H₂O₂ via the Fe²⁺-dependent, metal-catalyzed histidine oxidation (MCHO) mechanism, where a Fenton-like reaction leads to the irreversible oxidation of specific histidine residues, inactivating PerR and thereby relieving the repression of peroxide detoxification genes.

Beyond transcriptional regulation, increasing evidence underscores the critical role of small RNAs (sRNAs) in fine-tuning bacterial antioxidant defenses. Importantly, several sRNAs are found to mediate the trade-off between metal homeostasis and oxidative stress. For example, under iron deficiency, RyhB in *E. coli* and PrrF1/PrrF2 in *Pseudomonas aeruginosa* are induced, repressing the iron-dependent SodB to conserve iron, albeit at the cost of increased oxidative susceptibility (Argaman et al., 2012; Massé and Gottesman, 2002; Schu et al., 2015; Wilderman et al., 2004). Similarly, manganese deficiency triggers the expression of the sRNA RsaC in *Staphylococcus aureus*, which represses the synthesis of Mn-dependent SodA, prioritizing manganese allocation while increasing oxidative vulnerability (Lalaouna et al., 2019; McFarlane et al., 2025). Other sRNAs more directly regulate antioxidant enzymes; for instance, NfiS in *Pseudomonas stutzeri* stabilizes the mRNA of catalase KatB (Zhang et al., 2019), and OsiA/OsiR in *Deinococcus radiodurans* stabilize catalase transcripts (Gao et al., 2020). However, the mechanisms that integrate transcriptional and post-transcriptional layers to dynamically coordinate iron sparing and antioxidant defense remain largely unknown.

Mycobacteria are Gram-positive bacteria distinguished by their lipid-rich cell walls. This genus includes major human pathogens such as *Mycobacterium tuberculosis* and *M. abscessus*, along with environmental species such as the model organism *M. smegmatis* (Dartois and Dick, 2024; Kinsella et al., 2021; T et al., 2020; Tortoli, 2014). A key component of their oxidative defense is the bifunctional, iron-containing catalase-peroxidase, KatG. This enzyme not only decomposes peroxides but also activates the prodrug isoniazid (García-Marín et al., 2023; Liu, 2024; Wakamoto et al., 2013; Wang et al., 2025; Zhang et al., 1992). Transcription of *katG* is tightly regulated by the iron-responsive regulator FurA, a PerR-homolog (Lee et al., 2018; Zahrt et al., 2001). Under non-stress conditions, FurA dynamically represses *katG* and its own gene by binding to their promoter regions. This repression maintains basal levels of KatG sufficient for normal peroxide detoxification (Lee et al., 2018; Voskuil et al., 2011). Under peroxide stress, FurA is inactivated through the MCHO mechanism, thereby relieving its transcriptional repression and significantly increasing KatG production to counteract oxidative damage (Sala et al., 2003). This regulation places KatG expression at the nexus of iron homeostasis and oxidative stress management.

While sRNAs provide an essential layer of post-transcriptional control in mycobacterial stress responses (Arnvig and Young, 2009; Bar-Oz et al., 2023; Gerrick et al., 2018; Hnilicová et al., 2014), their specific roles in mediating adaptation to oxidative stress remain largely unexplored. In this study, we describe a coherent feed-forward loop that integrates transcriptional and post-transcriptional regulation to fine-tune KatG levels in response to iron and peroxide availability. We show that under iron-limiting conditions, the iron-sparing sRNA MrsI post-transcriptionally represses *katG1* mRNA, working in concert with the transcriptional repressor FurA3 to conserve iron. Conversely, under peroxide stress, an RNA sponge derived from the 3’-UTR of *furA3* is upregulated, sequesters MrsI, and thereby derepresses *katG1*, enabling rapid antioxidant defense. This multi-layered regulatory circuit allows mycobacteria to dynamically balance the competing demands of iron conservation and oxidative stress protection, revealing a sophisticated adaptive strategy for surviving in fluctuating environments.

## Results

### The sRNA MrsI significantly enhances the sensitivity of mycobacterial cells to oxidative stress

To investigate the regulatory roles of mycobacterial sRNAs in oxidative stress responses, we conducted sRNA sequencing to analyze intergenic sRNAs in *M. smegmatis* strain mc^2^155. By integrating our data with a published RNA-seq dataset (GEO: GSE62152) (Shell et al., 2015), we identified 40 sRNAs with precisely mapped 5’- and 3’-ends for further analysis (Appendix Table S1 and Appendix Fig. S1). We then overexpressed these sRNAs using the strong 16S rRNA promoter, P*_rrnB_*, and assessed their effects on the oxidative stress response. The results showed that while most sRNAs had no detectable effect, MrsI (ncMs13628Ac) markedly increased the susceptibility of mycobacterial cells to H₂O₂ (Fig. 1a). Given that the strong P*_rrnB_* promoter might amplify phenotypic effects, we switched MrsI overexpression to the moderate inducible P*_tetO_* promoter for finer control.

**Fig. 1.**
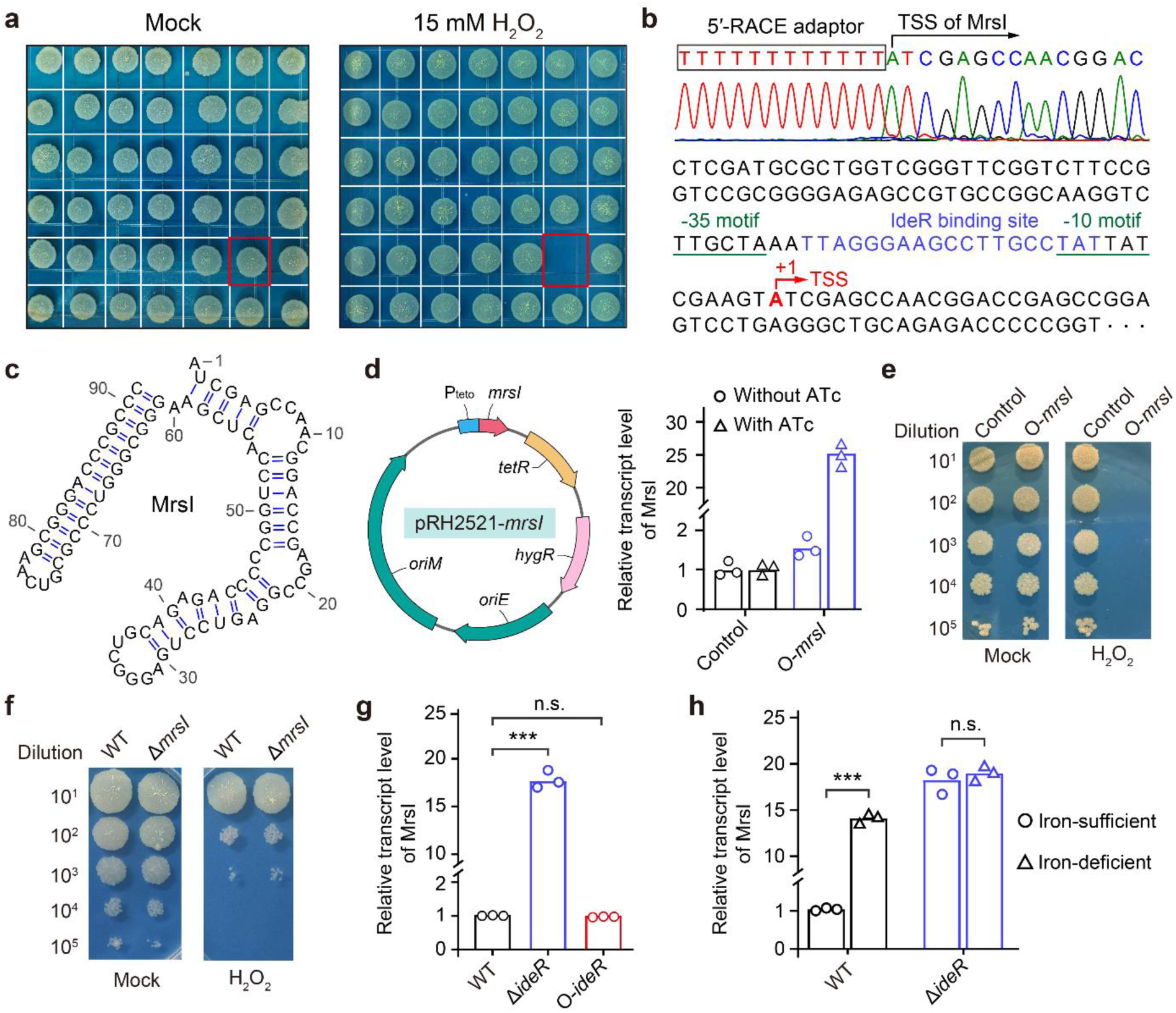
The sRNA MrsI enhances the sensitivity of mycobacterial cells to H₂O₂. **a**, H₂O₂ sensitivity of sRNA-overexpressing strains. The sRNA coding sequences were cloned into a multicopy plasmid under the control of the strong constitutive promoter P*_rrnB_*. Colonies boxed in red indicate strain overexpressing *mrsI*. **b**, Identification of the transcription start site (TSS) of MrsI by 5’-RACE. The IdeR binding site and TSS are marked in blue and red, respectively, while the core promoter elements are underscored. **c**, The secondary structure of *M. smegmatis* MrsI was predicted based on structural homology to the *M. tuberculosis* homolog. **d**, Construction of the ATc-inducible *mrsI* expression strain O-*mrsI*. The left figure shows the construction map of the overexpression plasmid; the right figure shows the MrsI transcript levels of strain with or without ATc induction. **e**, Sensitivity of the O-*mrsI* strain to H₂O₂. Mycobacterial cells were treated with 15 mM H₂O₂ for 1 h. **f**, Sensitivity of the *mrsI* deletion strain to H₂O₂. Mycobacterial cells were treated with 25 mM H₂O₂ for 1 h. **g**, qRT-PCR assay to measure MrsI transcript levels in *ideR* knockout and overexpression strains. **h**, qRT-PCR assay to measure MrsI transcript levels in the *ideR* knockout strain under different iron concentrations. Statistical analysis was done using the Student’s t-test. *** indicates *p*-value < 0.001, and n.s. indicates *p*-value > 0.05. **d**-**g,** The control strain is mc^2^155 harboring the empty pRH2521 vector. “Mock” indicates an untreated control samples.

To accurately locate the MrsI transcript, we used 5’-rapid amplification of cDNA ends (5’-RACE) to identify its 5’-end. As shown in Fig. 1b, an adenine (A) corresponding to position 3690371 in the mc^2^155 genome was identified at the 5’-end, along with canonical SigA-recognized -10 and -35 motifs upstream. This indicates that *mrsI* transcription is regulated by SigA and confirms that the mapped 5’-end corresponds to its transcription start site (TSS). Moreover, combining transcriptome data with predicted secondary structure analysis, we identified a stable hairpin structure at the 3’-end of MrsI (Fig. 1c). This feature is consistent with typical characteristics of mycobacterial transcription terminators (Hazra and Mukhopadhyay, 2025; Mitra et al., 2008), strongly suggesting that the hairpin structure mediates the transcriptional termination of the MrsI transcript. After confirming the full-length MrsI transcript, we placed its gene under the control of the ATc-inducible P*_teto_* promoter to construct the overexpression strain O-*mrsI* (Fig. 1d). Following ATc induction, the O-*mrsI* strain demonstrated a significant increase in MrsI levels along with a pronounced sensitivity to H₂O₂ (Fig. 1 d and 1e). These results are consistent with our previous observations and collectively demonstrate that MrsI plays a key role in the mycobacterial oxidative stress response.

To better elucidate the regulatory role of MrsI in the oxidative stress response, we constructed an *mrsI* deletion mutant (Δ*mrsI*) and assessed its antioxidant capacity. The Δ*mrsI* strain showed no significant difference in H₂O₂ sensitivity compared to the wild-type (WT) strain. (Fig. 1f). Consistent with this observation, we identified an iron-dependent regulator (IdeR) binding motif spanning the -10 to -35 region of the *mrsI* promoter (Fig. 1b), indicating that IdeR constitutively represses *mrsI* expression in the WT strain. This repression may explain why the Δ*mrsI* mutant remains sensitive to H₂O₂. To test this speculation, we quantified MrsI levels in *ideR* knockout (Δ*ideR*) and overexpression (O-*ideR*) strains. As shown in Fig. 1g, MrsI levels were significantly upregulated in the Δ*ideR* strain, confirming that IdeR indeed inhibits *mrsI* expression. Notably, IdeR overexpression did not further reduce MrsI levels (Fig. 1g), indicating that *mrsI* expression is already maximally repressed in the WT background. Given that IdeR activity is iron-dependent (Pandey and Rodriguez, 2014; Ranjan et al., 2006; Sritharan, 2016), we assessed the effects of iron availability on MrsI levels. Strikingly, MrsI levels were significantly upregulated under iron-deficient conditions (Fig. 1h), whereas this upregulation was abolished in the Δ*ideR* strain (Fig. 1h). Collectively, these results demonstrate that IdeR represses *mrsI* expression in an iron-dependent manner. Given that *mrsI* is highly inducible under diverse conditions (Gerrick et al., 2018), we prioritized the *mrsI* overexpression strain O-*mrsI* in subsequent experiments to accurately mimic its physiological expression pattern during stress conditions.

### MrsI directly targets *katG1* to regulate mycobacterial oxidative sensitivity

To elucidate the molecular basis of MrsI-mediated oxidative sensitivity, we first computationally predicted MrsI’s binding targets. Since sRNAs typically bind to the UTRs of target mRNAs, we extracted all 5’-UTR and 3’-UTR sequences from mc^2^155 and analyzed their potential interactions with MrsI. The results showed that 49 UTRs contained sequences that were fully complementary to the MrsI seed region (GGGCUG) (Gerrick et al., 2018). After careful analysis, we identified the catalase-peroxidase gene *katG1* (*MSMEG_6384*) as one of the predicted targets, with an interaction free energy of -11.35 kcal/mol, ranking relatively high among all predicted interactions (Appendix Table S2). To verify this interaction, we carried out an RNA-RNA electrophoretic mobility shift assay (EMSA). As shown in Fig. 2a, the MrsI band gradually retarded with increasing levels of the *katG1*_5’-UTR, indicating that the two RNA sequences formed a stable complex. When the key interacting nucleotides in the *katG1*_5’-UTR were mutated (*katG1*M), it failed to bind to MrsI and prevent its migration (Fig. 2b); similarly, when the key interacting nucleotides in MrsI were mutated (MrsIM), it also failed to bind to the *katG1*_5’-UTR and blocked its migration (Fig. 2c). Moreover, when we simultaneously mutated both RNA sequences to restore Watson-Crick pairing, they regained their binding ability and exhibited a retarded migration (Fig. 2d). These results indicate that MrsI can directly and specifically bind to *katG1*_5’-UTR.

**Fig. 2.**
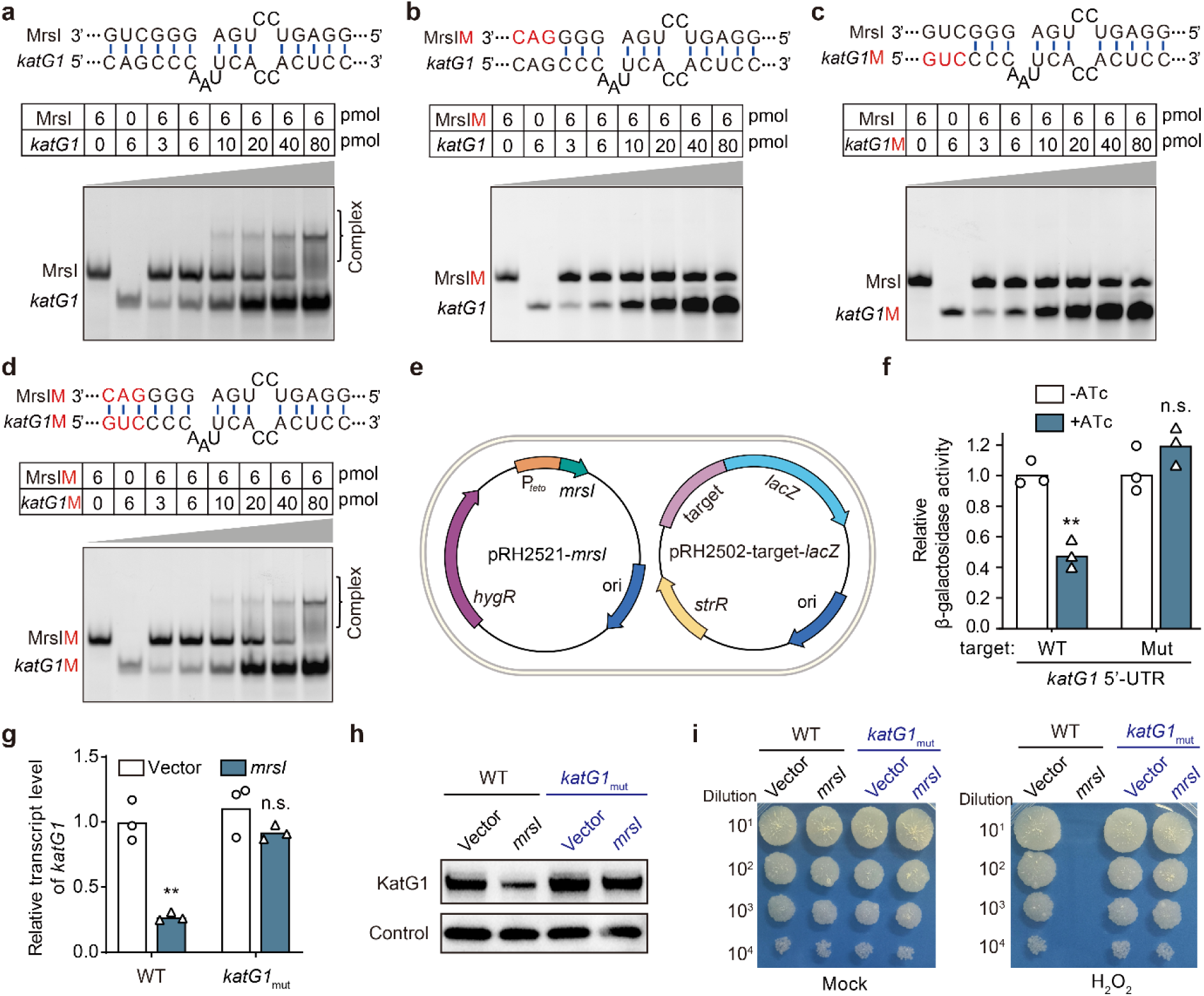
MrsI directly targets the 5’-UTR of *katG1*, inhibiting its expression and mediating oxidative sensitivity in mycobacterial cell. The interaction between MrsI and the *katG1*_5’-UTR was verified by RNA-RNA EMSA experiments. **a-d,** Interactions between WT MrsI and the WT 5’-UTR of *katG1*, WT MrsI and mutant 5’-UTR of *katG1* (*katG*M), mutant MrsI (MrsIM) and WT 5’-UTR of *katG1*, and mutant MrsI (MrsIM) and mutant 5’-UTR of *katG1* (*katG*M), respectively. **e,f,** The interaction between MrsI and *katG1* 5’-UTR was verified using a LacZ reporter system. **g**,**h**, KatG1 transcript and protein levels were measured in different strains using qRT-PCR and Western blot, respectively. The global transcription factor CarD was used as an internal control in Western blotting experiments. **i**, Effect of the *katG1*_5’-UTR sequence on MrsI-mediated regulation of oxidative sensitivity.

To further validate the interaction between MrsI and the *katG1* 5’-UTR, we conducted a LacZ reporter assay (Fig. 2e). As shown in Fig. 2f, after inducing MrsI expression by ATc, we observed a significant decrease in β-galactosidase activity in the strain carrying the WT *katG1*_5’-UTR (5’-UTR_WT_-*lacZ*), whereas no significant change was observed in the strains carrying the mutant *katG1*_5’-UTR (5’-UTR_Mut_-*lacZ*). This result is consistent with the EMSA data described above, confirming that MrsI directly bind to the 5’-UTR of *katG1*. Furthermore, we mutated the sequence within the 5’-UTR of *katG1* in the mc^2^155 genome (changing CAGCCC_6438880-6438885_ to CAaaCC_6438880-6438885_), constructed the mutant strain *katG1*_mut_, and assessed changes in MrsI-mediated regulation of *katG1*. As shown in Fig. 2g and h, MrsI effectively reduced KatG1 levels in the WT mc^2^155 strain but failed to achieve this effect in the *katG1*_mut_ strain. Furthermore, overexpression of *mrsI* significantly enhanced mycobacterial sensitivity to H₂O₂ in the WT strain, an effect that was significantly attenuated in the *katG1*_mut_ strain (Fig. 2i). Collectively, these results indicate that MrsI directly interacts with the 5’-UTR of *katG1* to regulate its expression, representing a major mechanism of MrsI-mediated regulation of mycobacterial oxidative sensitivity.

Beyond its role in oxidative detoxification, KatG1 can also activate the first-line anti-tuberculosis drug isoniazid by converting its prodrug form into a reactive free radical, thereby killing mycobacterial cells (Appendix Fig. S2). Therefore, we assessed the effect of MrsI on isoniazid resistance in mycobacterial cells. The results showed that MrsI significantly enhanced mycobacterial resistance to isoniazid but had no effect on other drugs (Appendix Fig. S3). Crucially, this effect was abolished in the *katG1*_mut_ strain (Appendix Fig. S3f), consistent with the pattern observed in oxidative stress regulation. Altogether, these findings confirm that MrsI downregulates *katG1* expression by binding to the 5’-UTR of *katG1*, thereby modulating oxidative stress sensitivity.

### MrsI directly targets the 3’-UTR of *furA3*

The expression of *katG* in mycobacteria is transcriptionally repressed by the ferric uptake regulator FurA, and there are three pairs of FurA and KatG in *M. smegmatis*, among which *furA1-katG1* and *furA2-katG2* are located in the same operons, while *furA3* and *katG3* are transcribed independently (Fig. 3a). Our results show that KatG1 is the main protein in mycobacterial cells to resist oxidative stress and activate isoniazid (Appendix Fig. S2), which is consistent with a previous report (Iwao and Nakata, 2018). Moreover, all three FurA proteins exhibit inhibitory effects on *katG1* expression (Lee et al., 2018). Notably, among all predicted MrsI targets, the 3’-UTR of *furA3* exhibits the highest affinity for MrsI, with an interaction free energy of -16.56 kcal/mol (Appendix Table S2). For convenience, this 3’-UTR is referred to as FutR. MrsI and FutR exhibit perfect complementarity of 10 base pairs (Fig. 3b), with the binding region located between the seed region of MrsI and the unstructured region of FutR (Fig. 3c and d). To verify this interaction, we carried out RNA-RNA EMSA. As shown in Fig. 3e, the MrsI band gradually retarded with increasing FutR concentration, indicating that the two RNA sequences formed a stable complex. When the key interacting nucleotides in FutR were mutated (FutRM), FutR was unable to bind to MrsI and blocked its migration (Fig. 3f); similarly, when the key interacting nucleotide in MrsI was mutated (MrsIM), FutR was also unable to bind to MrsI and prevent its migration (Fig. 3g). Moreover, when we simultaneously mutated these two RNA sequences to restore Watson-Crick pairing, they regained their ability to bind and exhibited a retarded migration (Fig. 3h). This indicates that MrsI can directly bind to FutR and that the binding between the two is specific.

**Fig. 3.**
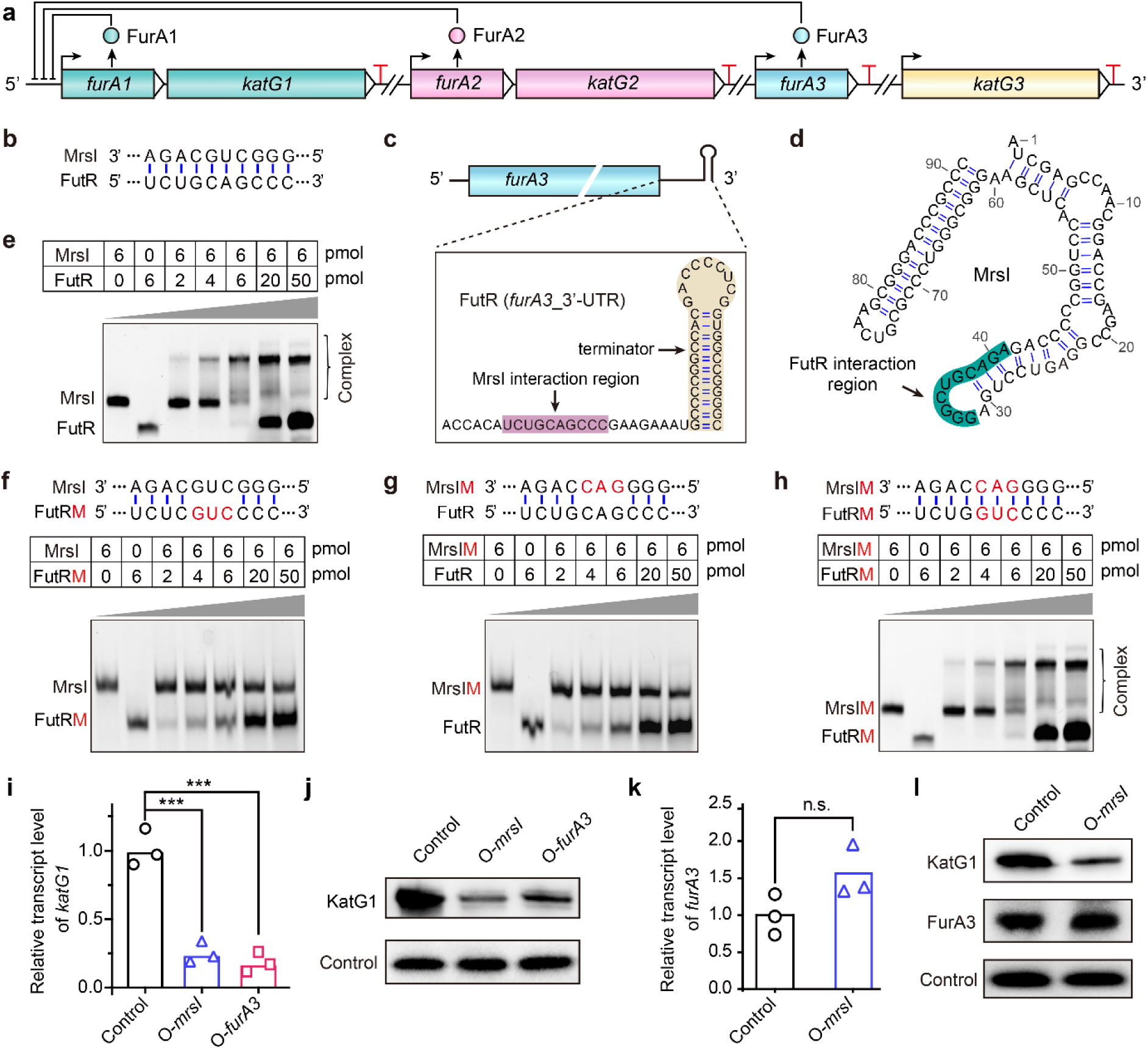
MrsI directly targets FutR in the 3’-UTR of *furA3*. **a**, Genomic location and regulatory network of *furA* and *katG* in *M. smegmatis*. Forward-curved arrow represents promoter, and red T-shaped symbol indicates the terminator. **b**, Potential interaction pattern between MrsI and FutR predicted by IntaRNA**. c**, Predicted secondary structure of FutR; the interaction region with MrsI is highlighted in pink, and the terminator region is highlighted in light yellow. **d**, Predicted secondary structure of MrsI, with the FutR interaction region highlighted in dark green. **e-h**, RNA-RNA EMSA experiments confirmed the interactions between WT MrsI and WT FutR, WT MrsI and mutant FutR (FutRM), mutant MrsI (MrsIM) and WT FutR, and mutant MrsI (MrsIM) and mutant FutR (FutRM), respectively. **i**,**j,** qRT-PCR and Western blot assays examined the regulation of KatG1 by MrsI and FurA3, respectively. **k**,**l,** qRT-PCR and Western blot assays examined the regulation of FurA3 by MrsI, respectively. CarD was used as an internal control in Western blot analysis. KatG1, FurA3, and CarD were detected using their respective polyclonal antibodies. Statistical test was done using the Student’s t-test, with *** indicating *p*-value < 0.001, and n.s. indicating *p*-value > 0.05. The control strain used in panels **i**-**l** was mc^2^155 harboring the empty pRH2521 vector.

Notably, when sRNAs bind to their target RNAs, their termini become spatially close and can be covalently linked by RNA ligases to form chimeric transcripts (Han and Lory, 2021; Nichols et al., 2008). This principle underlies the GRIL-seq technique, commonly used for sRNA target identification (Han et al., 2016). To further validate the interaction between MrsI and FutR, we expressed T4RNL1 (a T4 phage-derived RNA ligase) in an MrsI-overexpressing strain and performed whole-transcriptome deep sequencing. Our sequencing data revealed many MrsI-FutR chimeric transcripts (Appendix Fig. S4), which is consistent with the EMSA results described above. These findings collectively demonstrate that MrsI directly targets FutR.

Given that FurA3 plays a key role in *katG1* regulation and oxidative stress response (Fig. 3i-j) and that MrsI directly binds to its 3’-UTR (Fig. 3b-h), we investigated whether MrsI regulates *furA3* expression. The results showed that overexpression of MrsI did not significantly change FurA3 RNA or protein levels (Fig. 3k and l), indicating that MrsI does not regulate *furA3* expression. Furthermore, we investigated whether the MrsI-FutR interaction affects *katG1* expression and the oxidative stress response. To address this issue, we introduced a site-directed mutation into the FutR sequence of the mc²155 genome (mutating TCTGCAGCCC_6318217-6318212_ to TCTcgtGCCC_6318217-6318212_), generating the mutant strain FutR_mut_ (Appendix Fig. S5a-b). The results showed that even with this mutation in the FutR binding site, MrsI retained its ability to regulate *katG1* expression (Appendix Fig. S5c) and oxidative sensitivity (Appendix Fig. S5d-e). Similarly, this mutation also did not affect MrsI-mediated regulation of isoniazid resistance in mycobacterial cells (Appendix Fig. S5f). These findings demonstrate that MrsI-mediated regulation of KatG1 levels and associated phenotypes is independent of its interaction with FutR.

### FutR acts as an RNA sponge to alleviate the inhibitory effect of MrsI on *katG1*

Through the above experiments, we identified two targets of MrsI: *katG1* and FutR. MrsI appears to bind to *katG1* and suppress its expression, whereas the biological significance of the MrsI-FutR interaction remains unclear. Given that MrsI binds to FutR with significantly higher affinity than to *katG1* (-16.56 *vs* -11.35 kcal/mol) (Figs. 2a and 3d), we speculate that FutR can competitively bind MrsI to prevent its interaction with *katG1*. To address this speculation, we carried out a competitive EMSA assay. As expected, in the presence of equimolar amounts of FutR and *katG1*, MrsI preferential binds to FutR (Fig. 4a and b). Next, we tested whether high level of FutR affects MrsI-mediated regulation of *katG1 in vivo*. To achieve FutR overexpression, we constructed a recombinant MrsI-FutR co-expression strain (O-dual, Appendix Fig. S6) and examined changes in KatG1 levels. As shown in Fig. 4c and d, KatG1 transcript and protein levels were significantly reduced in the O-*mrsI* strain, while they were notably restored in the O-dual strain, indicating that high levels of FutR can relieve the inhibitory effect of MrsI on *katG1*. To confirm this observation, we examined the sensitivity of these strains to H₂O₂. The results showed that expressing MrsI alone significantly increased mycobacterial sensitivity to H₂O₂, whereas co-expression of MrsI and FutR nearly restored sensitivity to normal levels (Fig. 4e). Furthermore, we observed a similar effect on MrsI-mediated regulation of isoniazid resistance in mycobacterial cells (Appendix Fig. S7). These findings are consistent with the EMSA results described above, suggesting that FutR functions as a sponge (decoy target) to attenuate the inhibitory effects of MrsI on *katG1*.

**Fig. 4.**
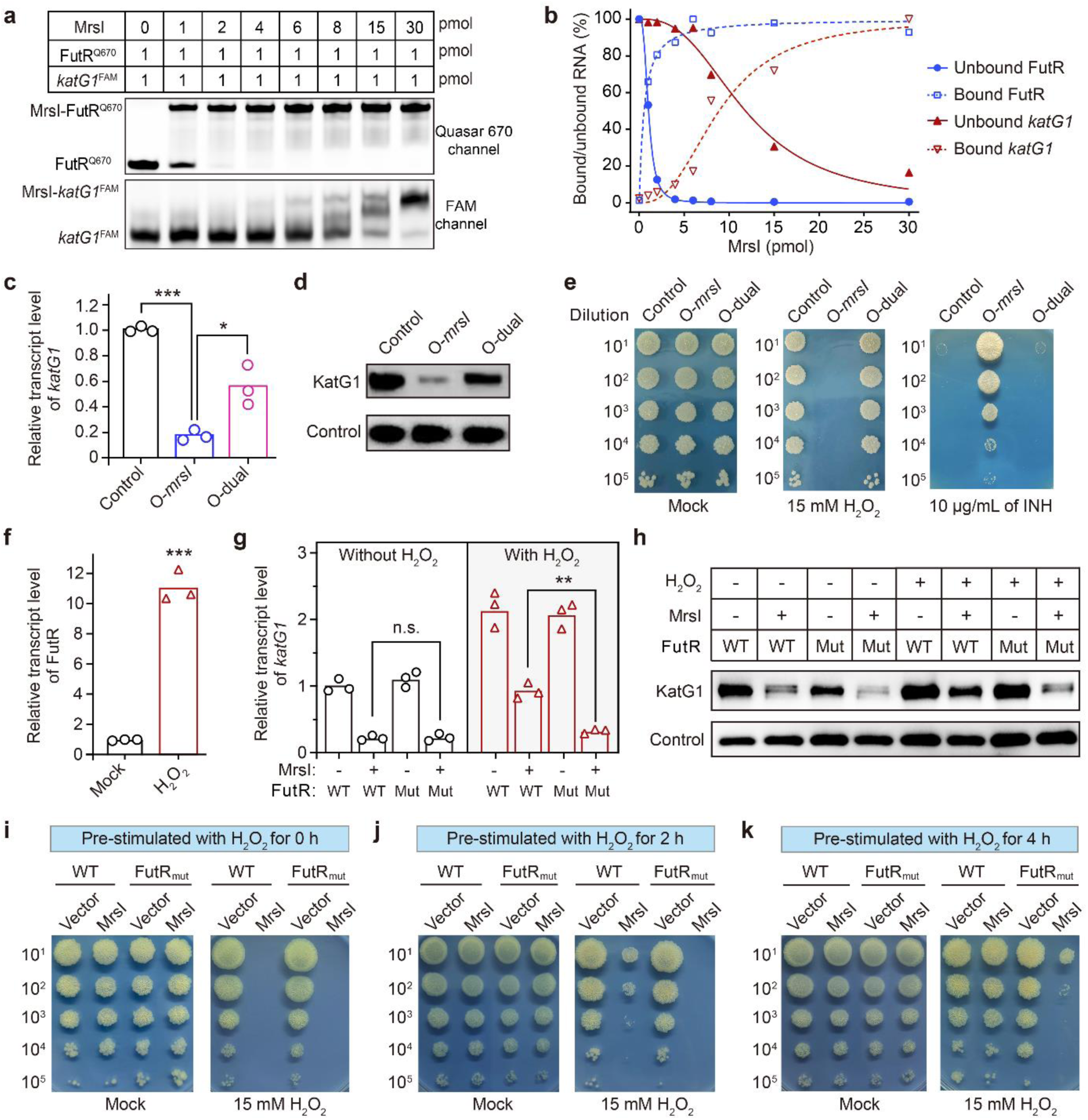
FutR acts as an RNA sponge to weaken the inhibitory effect of MrsI on *katG1*. **a**, Competitive RNA-RNA EMSA assays were used to examine the binding affinity of FutR and *katG1* to MrsI. FutR and *katG1* were labeled with Quasar 670 (Q670) and FAM, respectively. **b**, Quantification of MrsI-bound and unbound FutR/*katG1* in representative gels from panel **a**. **c,d**, qRT-PCR and Western blot assays to examine the transcript and protein levels of KatG1, respectively. **e**, Resistance plate assays were used to detect isoniazid resistance in O-*mrsI* and O-dual strains. **f**, qRT-PCR experiments were used to examine FutR transcript level in response to oxidative stress. **g**,**h**, qRT-PCR and Western blot assays were used to examine KatG1 transcript and protein levels, respectively. For oxidative stimulation experiments, mycobacterial cells were first grown to an OD_600_ of 0.5, followed by treatment with 0.5 mM H_2_O_2_ for 4 h. **i-k**, The role of FutR in mycobacterial cell resistance to oxidative stress. Mycobacterial cells were pre-stimulated with 0.5 mM H_2_O_2_ for 0 h (**i**), 2 h (**j**), and 4 h (**k)** before exposure to 15 mM H_2_O_2_. Statistical tests were done using Student’s t-test, with *** indicating *p*-value < 0.001, ** indicating *p*-value < 0.01, * indicating *p*-value<0.05, and n.s. indicating *p*-value > 0.05. For panels **c**-**e**, the control strain used is mc^2^155 harboring the empty pRH2521 vector.

It should be noted that the high level expression of FutR in the above experiments was achieved by artificial overexpression. However, can FutR expression reach a comparable level under physiological conditions? Considering that *furA3* is significantly induced under oxidative stress (Lee et al., 2018), we measured the level of FutR under oxidative challenge conditions. The result showed that FutR levels were robustly upregulated after H₂O₂ treatment (Fig. 4f), indicating that FutR acts as an RNA sponge under oxidative stress conditions. To mimic oxidative stress under physiological conditions, we used a low concentration (0.5 mM) of H₂O₂ to induce FutR instead of artificial overexpression. As shown in Fig. 4g-h, MrsI significantly suppressed *katG1* expression in both WT and FutR_mut_ strains without H₂O₂ pre-stimulation. However, after oxidative pre-stimulation, this inhibitory effect was attenuated in the WT strain but maintained in the FutR_mut_ strain. These results suggest that the induction of FutR under oxidative stress can counteract MrsI-mediated suppression of *katG1*, consistent with observations in FutR overexpression strain.

Furthermore, we investigated the regulatory role of FutR in mycobacterial oxidative detoxification. Critically, overexpression of MrsI in both the WT and FutR_mut_ strains resulted in high sensitivity to high-dose H₂O₂ treatment without H₂O₂ pre-stimulation (Fig. 4i). However, the WT strain overexpressing MrsI began to exhibit some antioxidant capacity after 2 hours of pre-stimulation and returned to normal antioxidant levels after 4 hours of pre-stimulation (Fig. 4j and k). In contrast, the FutR_mut_ strain overexpressing MrsI showed no antioxidant capacity after 2 hours of pre-stimulation and only weak antioxidant capacity after 4 hours of pre-stimulation (Fig. 4j and k). These results indicate that FutR is induced upon oxidative stress and that elevated FutR levels act as an RNA sponge to relieve the inhibitory effect of MrsI on *katG1*, thereby assisting mycobacterial cells in coping with oxidative stress.

### Regulation of KatG by MrsI appears to be conserved across mycobacteria

To assess the conservation of the KatG-MrsI-FutR regulatory system, we analyzed the distribution of the three components and their potential for complementary base-pairing across 115 mycobacterial reference genomes. Our analysis revealed that both *katG* and *mrsI* are each present in 99 species, with 96 species harboring both genes (Fig. 5a). Multiple sequence alignment and secondary structure analyses revealed that MrsI homologs share a highly conserved seed sequence (GGGCUG; Fig. 5b and S8), located within an accessible loop region (Fig. 5b), facilitating potential targeting interactions. Crucially, 89.6% (86 of 96) of the *katG* 5’-UTR were fully complementary to the MrsI seed region, indicating that MrsI-mediated *katG* regulation is highly conserved. To experimentally validate this regulation, we selected eight representative species for further investigation, including *M. tuberculosis*, *M. bovis*, *M. africanum*, *M. canettii*, *M. marinum*, *M. abscessus*, *M. intracellulare*, and *M. yongonense*. Because the high pathogenicity of most relevant mycobacterial species prevents their routine use in the general laboratory, we used a heterologous system in the non-pathogenic model organism *M. smegmatis*. We replaced its native *mrsI* gene and *katG* 5’-UTR sequences with orthologous sequences from these pathogenic strains (Appendix Table S3) to reconstruct the potential regulatory pair (Fig. 5c-g). Heterologous expression analysis revealed that in most tested strains, such as the pathogenic *M. abscessus* (Fig. 5d) and *M. intracellulare* (Fig. 5g), MrsI strongly inhibited the expression of its cognate *katG*, whereas in the *M. tuberculosis* group (Fig. 5e), the inhibitory effect was comparatively weaker. These findings indicate that MrsI-mediated *katG* regulation is widespread among mycobacterial species, although its extent varies among strains.

**Fig. 5.**
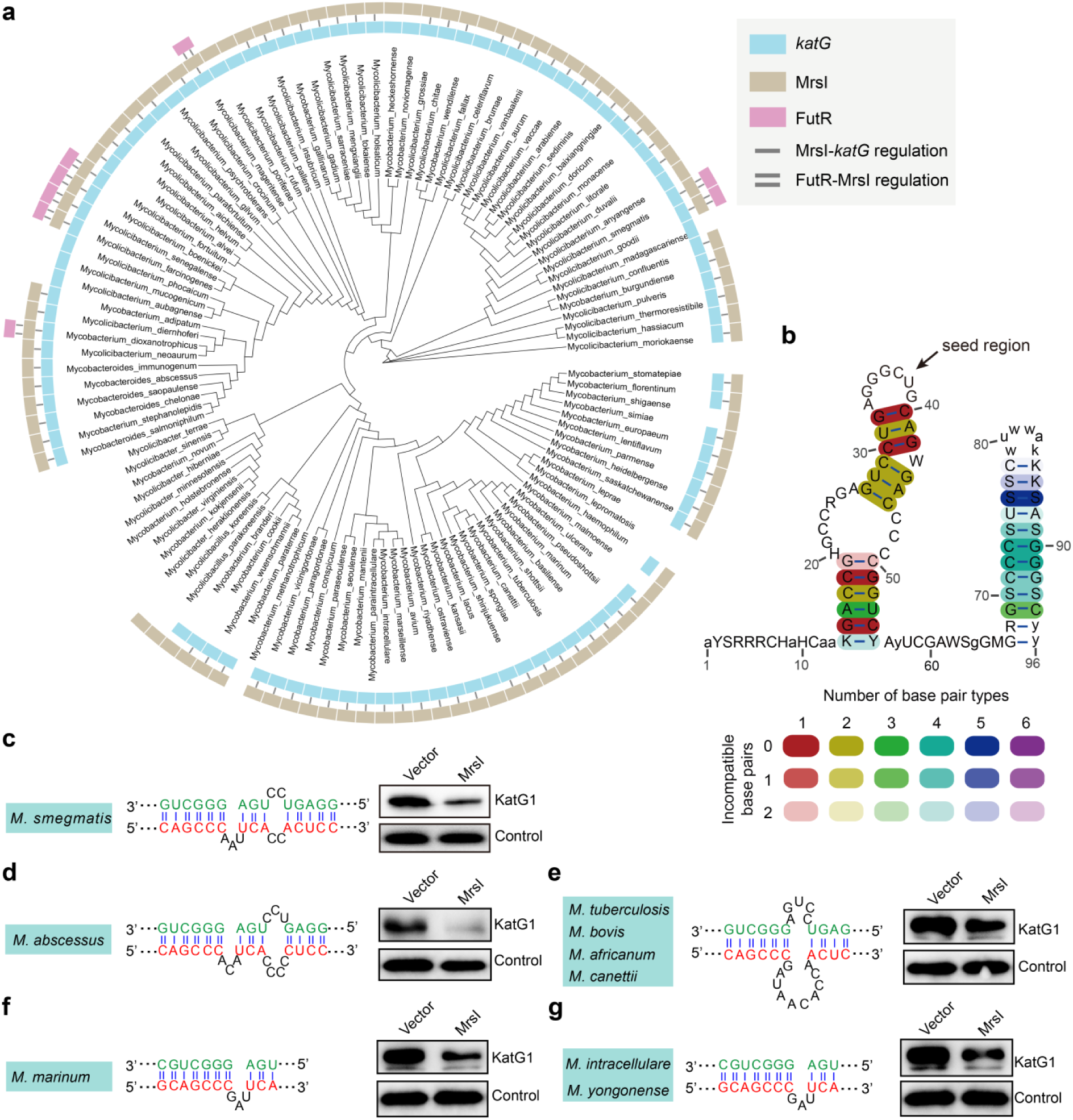
Conservation analysis of MrsI-*katG* regulation in mycobacteria. **a**, Distribution of *katG*, *mrsI*, and *futR* in 115 mycobacterial reference genomes. The phylogenetic tree was constructed based on the 16S rRNA sequences of these strains. The blue, brown, and pink rectangles indicate the presence of *katG*, *mrsI*, and *futR*, respectively, in the corresponding strains; blank spaces indicate the absence of the corresponding gene. The single line between the blue and brown rectangles indicates complementary pairing between MrsI and the *katG* 5’-UTR, and the double line between the brown and pink rectangles indicates complementary pairing between FutR and MrsI. **b**, Conservation of MrsI secondary structure. This structure was drawn using the LocARNA server. The arrow indicates the seed region. **c-g**, Validation of the conservation of the MrsI-*katG* regulatory mechanism across different mycobacterial species. Left: Schematic diagram of the predicted MrsI-*katG* base complementarity. Green nucleotides denote interacting residues on MrsI, and red nucleotides indicate corresponding residues on *katG* mRNA. Right: Western blot analysis of KatG protein levels in engineered bacterial strains. All immunoblotting experiments used CarD as an endogenous loading control. All experiments were carried out in triplicate, and representative images are shown. The MrsI and *katG*_5’-UTR sequences of *M. tuberculosis*, *M. bovis*, *M. africanum*, and *M. canettii* are identical (**e**), while those of *M. intracellulare* and *M. yongonense* are exactly the same (**g**).

Furthermore, we analyzed the distribution of FutR across these reference genomes. Our results revealed that FutR is present only in certain environmental mycobacterial species, such as *M. smegmatis*, *M. fortuitum*, and *M. goodie* (Fig. 5a). To comprehensively investigate the distribution and conservation of FutR, we expanded our analysis to encompass all sequenced mycobacterial genomes. FutR was identified in 40 strains from 14 species (Appendix Fig. S9), but its distribution was restricted to certain environmental species. In all 40 strains, FutR was located within the 3’-UTR of *furA*, indicating that its expression may also be induced by oxidative stress. Moreover, multiple sequence alignment revealed that FutR is relatively conserved, with 10 identical nucleotides critical for MrsI binding (Appendix Fig. S9), suggesting that its sponging function may be retained. Collectively, these data suggest that MrsI-mediated regulation of oxidative stress is a widely conserved mechanism in mycobacteria, while the FutR-mediated RNA sponging mechanism varies depending on the ecological niche.

## Discussion

Bacteria must constantly manage the fundamental trade-off between utilizing scarce iron for antioxidant defense and conserving this metal for essential cellular processes (Andrews et al., 2003; Imlay, 2013). In this study, we identify an extended coherent feed-forward loop in mycobacteria that integrates transcriptional and post-transcriptional regulation to dynamically balance iron homeostasis and oxidative stress defense. This circuit comprises the iron-responsive transcriptional regulator FurA3, the iron-sparing sRNA MrsI, its cognate RNA sponge FutR, and the iron-containing catalase-peroxidase KatG1 (Fig. 6). In the absence of external stimuli (Fig. 6 a), FurA3 binds to the promoter regions of *katG1* and its own gene (Lee et al., 2018), regulating *katG1* expression to maintain basal antioxidant capacity. Upon iron deficiency (Fig. 6b), MrsI is induced and binds to the 5’-UTR of *katG1* mRNA, significantly repressing its expression (Figs. 1 and 2). Importantly, when iron deficiency coincides with oxidative stress, mycobacteria upregulate KatG1 through a dual regulatory mechanism to counteract this threat (Fig. 6d). At the transcriptional level, FurA3 senses H₂O₂, undergoing conformational inactivation and relieving its repression of *katG1* and *futR* transcription (Figs. 3 and 4). At the post-transcriptional level, the induced RNA sponge FutR sequesters MrsI, relieving *katG1* mRNA repression and promoting the production of catalase-peroxidase, leading to efficient peroxide detoxification (Fig. 4). These findings reveal a multi-layered regulatory strategy that enables mycobacteria to fine-tune their antioxidant defense mechanism in response to fluctuating iron, highlighting the importance of post-transcriptional RNA networks in bacterial stress adaptation.

**Fig. 6.**
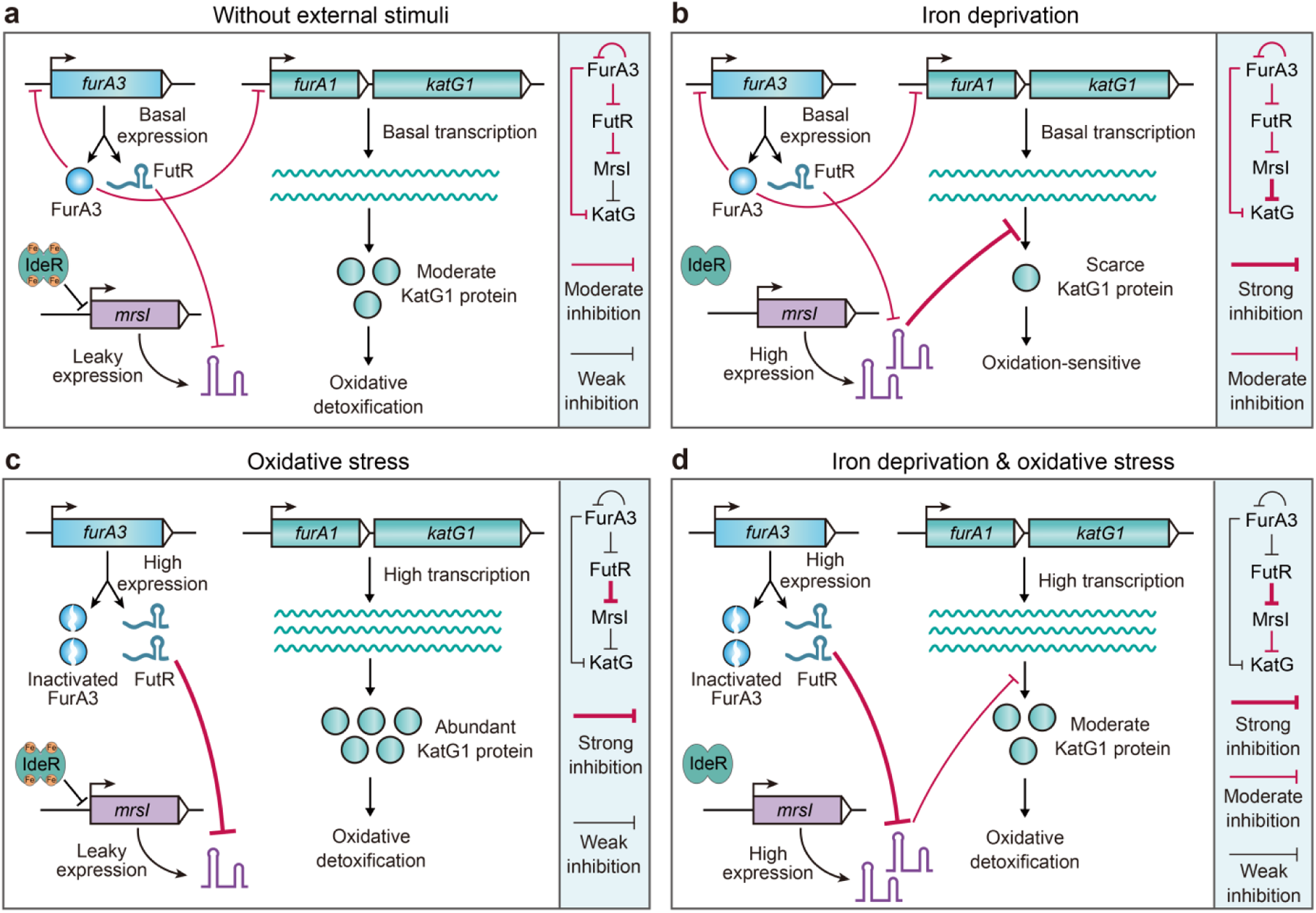
Model of a coherent FurA3-FutR-MrsI-KatG1 feed-forward loop for fine-tuning the oxidative stress response. **a,** Under basal conditions, the transcriptional regulator FurA3 binds to the promoter regions of *furA1*-*katG1* and its own gene (containing *futR*), dynamically regulating their expression and maintaining basal levels. Meanwhile, MrsI expression is repressed by IdeR. Basal expression of FutR sequesters leaky MrsI, preventing unnecessary metabolic disturbances. **b**, Under conditions of iron deficiency, IdeR is inactivated, leading to the release of MrsI expression. High levels of MrsI bind to the 5’-UTR of *katG1* mRNA, repressing its expression and increasing mycobacterial susceptibility to oxidative stress. **c**, Under oxidative stress, FurA3 is inactivated, thereby relieving the transcriptional repression of *katG1* and FutR. This leads to increased production of KatG1 and enhanced bacterial antioxidant capacity. **d**, Under the combined effects of iron deficiency and oxidative stress, FurA3 undergoes conformational inactivation and relieves its repression of *katG1* and *futR* transcription. In turn, induction of FutR, sequesters MrsI, thereby post-transcriptionally relieving the repression of *katG1* and promoting the rapid production of catalase-peroxidase, thereby achieving efficient detoxification. In each panel, the light blue box on the right depicts the regulatory relationships between the components.

MrsI, an iron-sparing sRNA, is a key mediator of iron conservation (Gerrick et al., 2018). Its repression of the iron-containing KatG1 represents a classic resource-allocation trade-off, conserving iron at the expense of antioxidant capacity. This strategy is analogous to that of *E. coli* RyhB, which represses the iron-dependent SodB under iron deficiency (Afonyushkin et al., 2005; Argaman et al., 2012; Massé and Gottesman, 2002). Notably, the activity of RyhB is modulated by the constitutively expressed RNA sponge 3’ETS*^leuZ^*, which primarily buffers transcriptional noise and fine-tunes the basal response (Lalaouna et al., 2015). In contrast, the FutR sponge identified here is specifically induced under oxidative stress in a FurA-dependent manner. This conditional induction allows FutR to function not merely as a noise filter, but as a critical switch that dynamically reprograms the post-transcriptional network in response to a concurrent stress, enabling a rapid shift from an iron-saving to an antioxidant-prioritizing state.

This conditional logic is central to the circuit’s function. Under oxidative stress, FurA inactivation and FutR induction form a coherent feed-forward loop that relieves *katG1* repression at both regulatory layers, driving robust KatG1 production. Once the peroxide threat is detoxified, FurA-mediated repression resumes, downregulating *futR* and releasing MrsI to restore iron-sparing regulation. Given that iron limitation is often a chronic challenge (Camaschella, 2019; Sun et al., 2024), whereas oxidative stress is typically acute (Buchser et al., 2023; Imlay, 2003, 2013), this circuit enables mycobacteria to dynamically prioritize the resolution of immediate, cytotoxic damage over persistent metabolic limitation, thereby optimizing fitness in fluctuating environments.

To our knowledge, FutR is the first reported RNA sponge in the Actinomycetota phylum. Phylogenetic analysis indicates that FutR is present in some environmental mycobacteria but absent in obligate intracellular pathogens like the *M. tuberculosis* complex (Fig. 5a). We propose that this distribution reflects distinct adaptive needs. Environmental species face highly variable oxidant levels (Imlay, 2019; Nguyen et al., 2021; Seixas et al., 2021), where the FutR-mediated, inducible switch provides a dynamic mechanism to rapidly recalibrate the iron-antioxidant trade-off in response to sudden stress. In contrast, obligate intracellular pathogens inhabit a more stable niche where host-derived oxidative stress is often persistent (Riffaud et al., 2023). In this context, the selective stress to maintain a rapid, condition-dependent post-transcriptional fine-tuner like FutR may be reduced, favoring reliance on direct transcriptional control. Thus, the presence or absence of FutR may signify an evolutionary adaptation to different lifestyles defined by the temporal dynamics of stress.

We also found that MrsI-mediated repression of KatG is broadly conserved across mycobacteria (Fig. 5a). Using *M. smegmatis* as a model, we experimentally validated this regulation in multiple pathogens, including the highly drug-resistant *M. abscessus* (Fig. 5d). While KatG suppression is a conserved response to iron deficiency, it may also contribute to the development of isoniazid resistance in patients. For instance, chronic malnutrition can lead to an anemia and an iron-deficient host environment (Nienaber et al., 2023), which may increase MrsI expression and downregulating KatG, thereby conferring isoniazid resistance. While drug resistance is often attributed to mutations (Farhat et al., 2024; Pei et al., 2024), our studies have uncovered a form of phenotypic resistance mediated by sRNA. This finding explains how mycobacterial cells resist antibiotic stress before acquiring mutations, providing new insights into the early stages of drug resistance.

## Methods

### Bacterial strains and growth condition

*E. coli* strains were cultivated in lysogeny broth (LB) medium at 37°C. *M. smegmatis* mc^2^155 WT strain and its derivatives were grown at 37°C in Middlebrook 7H9 medium supplemented with 0.5% (v/v) glycerol and 0.05% (v/v) Tween 80, or on Middlebrook 7H10 agar supplemented with 0.5% (v/v) glycerol. When needed, antibiotics were added at the following concentrations: kanamycin sulfate (Kan), 25 μg/mL; hygromycin B (Hyg), 50 μg/mL; streptomycin sulfate (Str), 10 μg/mL. The key bacterial strains used in this study are provided in Appendix Table S4.

### Construction of sRNA overexpression strains

sRNA overexpression strains were constructed based on multi-copy plasmid pMV261. Specifically, the DNA sequences of sRNA were cloned into the BamHI and HindIII restriction site of the pMV261 vector, which enabled expression of the sRNAs by the P*_rrnB_* promoter. For the construction of a strain that can inducibly express MrsI, the pRH2521 plasmid was used (Singh et al., 2016). Specifically, the *mrsI* DNA sequence was fused to the P*_teto_* promoter using an overlapping PCR strategy. This design enables the P*_teto_* promoter to regulate *mrsI*, thereby promoting efficient induction upon the addition of anhydrotetracycline (ATc). The pRH2521 plasmid was linearized with ClaI enzyme and then ligated with the P*_teto_*-*mrsI* fragment using Gibson assembly. The overexpression plasmid was then transformed into mc^2^155 cells to obtain an overexpression strain. The primers used are listed in Appendix Table S5.

### CRISPR/Cpf1-mediated mutagenesis

CRISPR/Cpf1-mediated mutagenesis of mycobacterial genes was carried out as previously described (Yan et al., 2017). This gene editing system includes three main parts: plasmid pJV53-cpf1, which expresses Cpf1 protein and RecE/T recombinase; plasmid pCR-Hyg that expresses the target gene crRNA; and a 59-nt single-stranded DNA targeting the lagging strand for homologous recombination repair. Initially, pJV53-cpf1 was electro-transformed into mc^2^155 cells to generate strain Ms/pJV53-cpf1. Once the OD_600_ of Ms/pJV53-cpf1 cells reached 0.5, 0.2% acetamide was introduced to induce the expression of RecET recombinase. Three hours later, the cells were harvested and prepared into competent cells for subsequent gene editing. Then, the plasmid pCR-Hyg containing the target gene crRNA and 59-nt ssDNA was electro-transduced into Ms/pJV53-cpf1 cells, and then spread into 7H10 resistance plates containing 25 μg/mL of Kan, 50 μg/mL of Hyg, and 50 ng/mL of ATc. After culturing at 37°C for 4 days, transformants were selected for PCR verification, and positive clones were further confirmed by sequencing. To eliminate plasmid, mutant strains were inoculated into antibiotic-free 7H9 medium and cultivated at 37°C. After 3-5 consecutive passages, strains were streaked onto antibiotic-free 7H10 plates. After colony growth, each colony was picked, and plasmid elimination was finally confirmed by PCR.

### β-Galactosidase reporter assay

The β-galactosidase reporter gene assay was carried out to validate the MrsI target. For exploring the interaction between MrsI and *katG1*_5’-UTR, the *mrsI* DNA sequence was inserted into the pRH2521 plasmid under the control of the P*_teto_* promoter and induced with 50 ng/mL of ATc. The *katG1*_5’-UTR and its upstream expression region (genome position: 6436771-6439111) were then cloned into the pRH2502 vector upstream of the *lacZ* gene and expressed under its endogenous promoter. Subsequently, both plasmids were transformed into the mc^2^155 strain to measure β-galactosidase activity. During the β-galactosidase activity assay, cells equivalent to an OD_600_ of 1 were harvested at specified intervals (e.g., 2 mL of each culture with OD_600_ = 0.5) and washed with 10 mM phosphate buffered saline (pH 7.4). Detailed procedures for β-galactosidase activity determination were conducted according to a previous reference (Tang et al., 2014).

### Identification of transcription start sites by 5’-RACE

To identify the TSS of MrsI, 5’-RACE analysis was done with RNA extracted from mc^2^155 cells grown in 7H9 medium. The 5’-RACE experiment was carried out as previously described (Zaunbrecher et al., 2009). The primers used are listed in Appendix Table S5.

### Measurement of protein levels by Western blot

For Western blot, the target proteins KatG1 and FurA3, along with the internal control protein CarD, were detected using their respective rabbit polyclonal antibodies. All experiments were conducted with three replicates, and a representative result is presented in the figure. Detailed procedures followed a previously described method (Li et al., 2022).

### Antibiotic stress survival assay

Mycobacterial cells were grown to mid-exponential phase (OD_600_≈1.0) and then diluted 10^1^, 10^2^, 10^3^, 10^4^, and 10^5^ folds, respectively. Afterward, 3 μL of bacterial suspension at each dilution level was spotted onto 7H10 plates containing isoniazid at different concentrations. Plates were cultivated at 37°C for 3 days before image collection.

### Measurement of transcript levels by qRT-PCR

Total RNA was extracted by the phenol/chloroform method as described previously (Li et al., 2017). The quality and concentration of total RNA were analyzed using NanoDrop 2000 (Thermo Scientific, Waltham, MA, USA). The reverse transcription experiment was conducted using a commercially available PrimeScript RT reagent kit with gDNA eraser (Takara Biotechnology, Tokyo, Japan) according to the manufacturer’s instructions. For qRT-PCR, *sigA* was used as the internal reference gene and the experiment was carried out according to the reference (Tang et al., 2014). The primers used in this study are listed in Appendix Table S5.

### RNA sequencing and sRNA identification

mc^2^155 cells equivalent to 30 OD_600_ (e.g., 30 mL of 1 OD_600_ of one culture) were harvested from mid-exponential, early-stationary and mid-stationary phases, respectively. Total RNA was extracted by the phenol/chloroform method as described previously (Li et al., 2017). For sRNA sequencing, 5 μg of total RNA from each sample was initially treated with RNase-free DNase I (Takara, Tokyo, Japan) to prevent potential genomic DNA contamination. Subsequently, ribosomal RNAs from Gram-positive organisms were removed using the RiboZero rRNA removal kit (Epicentre, Madison, WI, USA) prior to sequencing analysis. A total of 100 ng of rRNA-depleted RNA from each sample was fragmented into 30 to 100 nucleotides and used as a template for randomly primed PCR. Strand-specific cDNA libraries were then generated using standard protocols for subsequent Illumina sequencing. The resulting cDNAs were sequenced on an Illumina HiSeq 2500 platform (San Diego, CA, USA). RNA-seq data have been submitted to SRA under the accession number PRJNA1105428.

The quality of the raw sequence data was first assessed using FastQC (v0.11.9), and the sequences were further filtered and trimmed with Trimmomatic (v0.39) (Bolger et al., 2014). Clean reads were then mapped to the reference genome of mc^2^155 (NC_008596.1) using BLASTN to determine their genomic locations. Specifically, during the BLASTN sequence alignment, each read was allowed to match only one position in the genome, with an E-value set to 1e-5. In the output, hits with a read coverage of less than 80% or a sequence identity of less than 100% were discarded. To obtain a single-nucleotide resolution transcriptome map, we used a custom-written Perl script to calculate the transcription levels of each nucleotide and visualized the transcription data using Artemis software (Carver et al., 2012). For sRNA identification, we manually inspected the entire genomic transcription profile with Artemis-based visualization. We selected intergenic-derived sRNAs based on the following three criteria: 1) obvious transcription signals in intergenic regions; 2) clear transcription start and termination signals in intergenic regions; 3) transcript lengths between 40 and 200 nt.

### Prediction of MrsI regulatory targets using IntaRNA

In predicting MrsI targets, we first extracted all the 5’-UTR and 3’-UTR sequences from *M. smegmatis* mc^2^155 and analyzed their potential interactions with MrsI. Specifically, in our previous study, we performed a global analysis of the TSS and transcripts of mc^2^155 (Li et al., 2017). Based on this data, we extracted the sequences between the TSS and the start codon to use as 5’-UTR sequences, and the 20 nt downstream of the last stop codon of the transcript as 3’-UTR (for monocistronic genes, this corresponds to the stop codon of the gene, and for polycistronic genes, it refers to the stop codon of the last gene in the transcript). For target prediction, IntaRNA was used with the seed-defined model (Mann et al., 2017).

### RNA-RNA electrophoretic mobility shift assay

The RNA-RNA EMSA was used for investigating the *in vitro* interactions between the sRNA MrsI and its targets, including the *katG1* 5’-UTR and *furA3* 3’-UTR. Unlabeled transcripts were used for individual binding assays and FAM-labeled *katG1* 5’-UTR (3’-end) and Quasar 670 labeled FutR (3’-end) were used for competitive experiments, as detailed in Appendix Table S5 for all transcript and mutant sequences. Reaction mixtures (20 μL total volume) containing specified concentrations of sRNA and target RNA were prepared in a buffer composed of 20 mM Tris-HCl (pH 8.0), 1 mM DTT, 1 mM MgCl₂, 20 mM KCl, and 10 mM Na₂HPO₄-NaH₂PO₄, followed by an initial 2-minute denaturation at 90°C to eliminate secondary structures. Following incubation at 37°C for 30 minutes to promote RNA-RNA complex formation, the samples were resolved through 8% TBE gel electrophoresis. Visualization was achieved using GoldView™ staining for unlabeled probes and FAM/Quasar 670 fluorescence imaging for labeled probes.

### Identification of MrsI-FutR chimeric transcripts by high-throughput sequencing

To validate the *in vivo* interaction between MrsI and FutR, a GRIL-seq-like approach was employed. The codon-optimized T4RNL1-encoding gene was cloned into the pRH2521 vector under the control of the P*_teto_* promoter, while the *E. coli* Hfq-encoding gene and the native promoter-driven MrsI-encoding gene were separately cloned into the pMV261 vector. Both plasmids were co-electrotransformed into mc^2^155 cells to generate the recombinant strain Ms/*T4RNL1_hfq_mrsI*. The recombinant strain was cultured at 37°C in Middlebrook 7H9 medium. When the culture reached an OD_600_ of 0.5, 50 ng/mL ATc was added to induce T4RNL1 expression. Cells were harvested 4 h post-induction by centrifugation (12,000×*g*, 2 min, 4°C). Total RNA was extracted using TRIzol reagent, and ribosomal RNA was depleted prior to RNA sequencing. Library preparation and paired-end sequencing were performed as previously described (Li et al., 2017). Raw sequencing data were deposited in the NCBI Sequence Read Archive under accession number PRJNA1105428.

### Secondary structure and conservation analysis of MrsI

The RNAfold (Lorenz et al., 2011) and LocARNA (Will et al., 2012) web servers were used to predict the secondary structure of MrsI and the conservation of its secondary structure, respectively. Additionally, the VARNA tool (Darty et al., 2009) was used to display and edit RNA secondary structures. To predict targets regulated by MrsI, the sRNARFTarget (Naskulwar and Peña-Castillo, 2022) and IntaRNA (Mann et al., 2017) web servers were used.

### Oxidative stress experiment

For the oxidative stress experiment, mycobacterial cells were first cultured in 7H9 medium containing 50 ng/mL of ATc to the early exponential phase (OD_600_ ≈ 0.5). The bacterial suspension was centrifuged at 6000×*g* for 2 minutes, after which the supernatant was removed and the bacterial pellets were resuspended in an equal volume of fresh ATc-free 7H9 medium. To induce FutR expression, a low concentration (0.5 mM) of H₂O₂ was added to the medium (Note: to prevent the complete decomposition of H₂O₂, fresh 0.5 mM H₂O₂ was added every hour), and at 0, 2, and 4 hours of this stimulation, 15 mM H₂O₂ was added to the mixture to assess the oxidative detoxification ability. The mixture was shaken at 37°C for 1 hour, followed by centrifuged at 6000×*g* for 2 minutes. Then, the supernatants were removed and the bacterial pellets were resuspended in an equal volume of fresh H₂O₂-free 7H9 medium. Finally, the mycobacterial cells were diluted 10¹, 10², 10³, 10⁴, and 10⁵-fold, and 3 μL of each dilution was spotted onto 7H10 plates, followed by incubation at 37°C for 3 days.

## Statistical analysis

Statistical analyses were performed using a two-tailed Student’s t-test. Data are represented as the mean ± SD (n=3 biological replicates, which are independent experiments conducted on distinct cultures). Significance levels are denoted as follows: *p*-value < 0.05, *p*-value < 0.01, *p*-value < 0.001; n.s. (not significant) corresponds to *p*-value > 0.05.

## Data availability

The data underlying this article are available in the article and in its online supplementary material.

## Supplemental information

Supplemental materials including: Appendix Figs. 1–9 and Appendix Table S1-5.

## Author contributions

Conceptualization, X.F.L., J.H. and H.Y.; Methodology, Y.Z.H., F.C. and X.Y.L.; Formal analysis, F.C., X.Y.T., and Y.Z.H.; Investigation, X.F.L., Y.Z.H., J.F.X. and X.Y.L.; Writing-Original Draft, X.F.L. and J.H.; Writing-Review & Editing, H.Y and J.H.; Visualization, Y.Z.H. and M.Y.Z.; Funding Acquisition, J.H. and X.F.L.; Supervision, J.H. and H.Y.

## Funding

This work was supported by the National Natural Science Foundation of China (grants 32200026, 32570179, and 32500128) and the China Postdoctoral Science Foundation (grant 2019M662654).

## Competing interests

The authors declare no competing interests.

